# DNA Methylation Drives Aberrant Osteochondrogenesis in Keloid and Is Reversible by Decitabine

**DOI:** 10.64898/2026.08.26.747276

**Authors:** Lian Zhang, Chenmei Liu, Xinyuan Zhou, Yan Zhang, Renliang He, Ziyan Li, Shuqing Zhao, Chengcheng Deng, Hong-Tao Li, Bin Yang

## Abstract

Keloids are benign fibroproliferative disorders majorly characterized by excessive extracellular matrix deposition, with recurrence rates exceeding 80% following conventional therapy. Although epigenetic dysregulation has been implicated in keloid pathogenesis, whether genome-wide DNA methylation actively drives pathological cellular reprogramming, and whether this state is therapeutically reversible, remains unclear. We performed genome-wide DNA methylation profiling on keloid tissues, matched primary keloid fibroblasts, and normal controls. Our analysis revealed a shared DNA hypermethylation pattern between keloid tissues and fibroblasts, which was validated by three independent public cohorts. By integrating DNA methylome and transcriptome, we demonstrated that DNA methylation-regulated genes were enriched in osteochondrogenesis-related pathways, such as cartilage and bone development pathways. Furthermore, pharmacologic inhibition of DNA hypermethylation by DNA demethylating agent decitabine reduced the expression of osteochondrogenic markers and inhibited collagen deposition and keloid growth in primary keloid fibroblasts and patient-derived xenograft (PDX) model, offering a potential therapeutic strategy of keloid.

## Introduction

Keloids are benign, tumor-like fibroproliferative disorders majorly characterized by pathological fibroblast hyperproliferation and aberrant extracellular matrix deposition^1,2^. Current first-line treatments, including intralesional corticosteroids, radiotherapy, and surgical excision, remain inadequate, with recurrence rates exceeding 80%^3,4^, underscoring the urgent need for mechanism-based therapeutic strategies.

A clinically distinctive feature of keloids is their predisposition to ectopic osteochondrogenic differentiation, manifested by cartilage- and bone-like molecular and histological features^5,6^. This aberrant differentiation reprogramming is driven in part by hyperactivation of BMP and WNT developmental signaling pathways^7,8^ and is strongly associated with excessive growth and high frequent recurrence, suggesting that keloids undergo pathological cellular reprogramming rather than simple fibroblast hyperproliferation.

Epigenetic regulation, particularly DNA methylation, plays a central role in stabilizing cell identity and differentiation states^9–11^. Unlike genetic mutations, DNA methylation is pharmacologically reversible, making it an attractive therapeutic target. Genome-wide studies have reported extensive DNA methylation alterations in keloids^12–14^, predominantly hypermethylation affecting tumorigenesis- and fibrosis-associated pathways^15,16^. However, two critical gaps remain. First, whether the observed methylation changes actively drive, rather than merely correlate with, the osteochondrogenic phenotype has not been established at the mechanistic level. Second, the genomic context (promoter versus gene-body) in which methylation exerts its dominant regulatory effect in keloids is unknown; gene-body hypermethylation has been shown to activate rather than silence transcription in cancer and may represent an underappreciated mechanism in fibrotic disease.

Decitabine (5-aza-2′-deoxycytidine), an FDA-approved DNA methyltransferase inhibitor, reverses pathological DNA methylation states in hematological malignancies^17,18^ and emerging evidence suggests that DNMT inhibition can modulate fibroblast differentiation and fibrotic remodelling^19–21^. Nevertheless, whether targeting DNA methylation can reverse the osteochondrogenic differentiation in keloids has not been investigated.

Here, we integrate genome-wide DNA methylation profiling and transcriptomic profiling of keloid tissues and primary keloid fibroblasts with cross validation in four independent public cohorts to show that aberrant DNA methylation is a key mechanism driving osteochondrogenic transcriptional activation in keloids. We further demonstrate that this pathological epigenetic state is reproduced in primary fibroblasts and is pharmacologically reversible by decitabine both in vitro and in a patient-derived xenograft model in vivo.

## Materials and Methods

### Ethics approval and consent to participate

All human subjects research conducted in this study was reviewed and approved by the Human Ethics Committee of the Dermatology Hospital of Southern Medical University. Written informed consent was obtained from patients. These experimental methods comply with Helsinki Declaration.

### Sample collection

Keloid specimens (n=14) were obtained from 4-mm punch biopsies taken during therapeutic procedures at the Dermatology Hospital of Southern Medical University. Control tissues (n=4) were acquired from surgery discards, with anatomical matching attempted when possible.

For RNA sequencing, fresh tissues were immediately stabilized in RNAlater™ Solution (Thermo Fisher, Cat#AM7021) at 4°C for 24h before -80°C storage. For DNA methylation analysis, matched specimens were snap-frozen in liquid nitrogen within 2 minutes post-excision and maintained at -80°C until processing.

Patients’ information was shown in table 1.

**Table 1.**
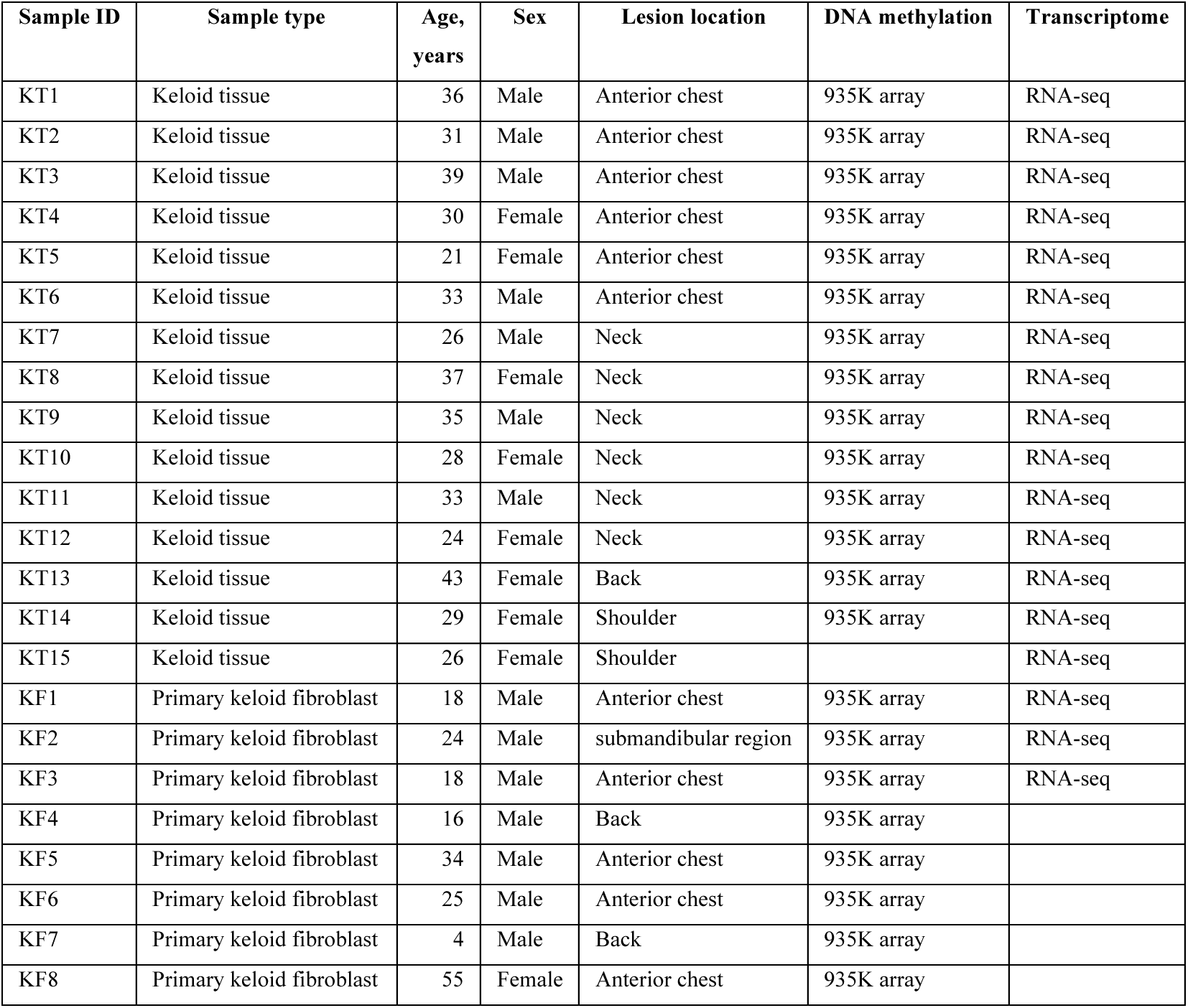
Clinical characteristics of the keloid tissue sample used for sequencing.

| Sample ID | Sample type | Age, years | Sex | Lesion location | DNA methylation | Transcriptome |
| --- | --- | --- | --- | --- | --- | --- |
| KT1 | Keloid tissue | 36 | Male | Anterior chest | 935K array | RNA-seq |
| KT2 | Keloid tissue | 31 | Male | Anterior chest | 935K array | RNA-seq |
| KT3 | Keloid tissue | 39 | Male | Anterior chest | 935K array | RNA-seq |
| KT4 | Keloid tissue | 30 | Female | Anterior chest | 935K array | RNA-seq |
| KT5 | Keloid tissue | 21 | Female | Anterior chest | 935K array | RNA-seq |
| KT6 | Keloid tissue | 33 | Male | Anterior chest | 935K array | RNA-seq |
| KT7 | Keloid tissue | 26 | Male | Neck | 935K array | RNA-seq |
| KT8 | Keloid tissue | 37 | Female | Neck | 935K array | RNA-seq |
| KT9 | Keloid tissue | 35 | Male | Neck | 935K array | RNA-seq |
| KT10 | Keloid tissue | 28 | Female | Neck | 935K array | RNA-seq |
| KT11 | Keloid tissue | 33 | Male | Neck | 935K array | RNA-seq |
| KT12 | Keloid tissue | 24 | Female | Neck | 935K array | RNA-seq |
| KT13 | Keloid tissue | 43 | Female | Back | 935K array | RNA-seq |
| KT14 | Keloid tissue | 29 | Female | Shoulder | 935K array | RNA-seq |
| KT15 | Keloid tissue | 26 | Female | Shoulder |  | RNA-seq |
| KF1 | Primary keloid fibroblast | 18 | Male | Anterior chest | 935K array | RNA-seq |
| KF2 | Primary keloid fibroblast | 24 | Male | submandibular region | 935K array | RNA-seq |
| KF3 | Primary keloid fibroblast | 18 | Male | Anterior chest | 935K array | RNA-seq |
| KF4 | Primary keloid fibroblast | 16 | Male | Back | 935K array |  |
| KF5 | Primary keloid fibroblast | 34 | Male | Anterior chest | 935K array |  |
| KF6 | Primary keloid fibroblast | 25 | Male | Anterior chest | 935K array |  |
| KF7 | Primary keloid fibroblast | 4 | Male | Back | 935K array |  |
| KF8 | Primary keloid fibroblast | 55 | Female | Anterior chest | 935K array |  |

### DNA extraction and EPIC DNA methylation data generation

Genomic DNA from keloid tissue and fibroblast cell were extracted using QIAamp DNA Mini Kit (QIAGEN, Cat#51304) according to the manufacturer’s introduction. 1 μg DNAs were then subject to bisulfite conversion using EZ DNA Methylation-Lightning Kit (Zymo Research, Cat#D5030) as described by the manufacturer. DNA methylation was evaluated using Illumina Infinium MethylationEPIC v2.0 BeadChip as recommended by the manufacturer. Briefly, the bisulfite converted DNA was amplified, enzymatically fragmented and hybridized overnight to EPIC v2.0 BeadChip, which was designed to detect the methylation status of over 935,000 CpG sites on human genome. BeadArrays were scanned using the Illumina iScan system to generate the IDAT files in both red and green channels. IDAT files were then processed by SeSAMe R package to generate beta (β) value of each CpG site by specifying collapseToPfx=TRUE to average the beta values of replicate probes sharing the same probe name prefixes. Beta value represents the DNA methylation value of each CpG site and ranges from 0 to 1, with zero for unmethylated sites and one for fully methylated sites.

### DNA methylation data analysis

The probes with detection *p* value greater than 0.05 as well as those linked with single nucleotide polymorphism (SNP), gender and age were removed from further analysis. We also removed the probes with missing value in any sample. Probes with absolute difference of β value (|Δβ|) > 0.2 between keloid and control groups were selected. Welch’s *t*-test was performed to identify significant difference of the methylation sites (p value < 0.05). Hierarchical clustering analysis and heatmap were generated using the R package ComplexHeatmap (v2.20.0). Principal component analysis was performed using the R function prcomp. Gene oncology analysis and functional annotation were performed using R package clusterProfiler (v4.12.6).

### Probe annotations

Probe annotations were obtained from the Infinium MethylationEPIC v2.0 manifest (hg38, illumina.com). We defined probe sites as (1) “Promoter” probes for those linked to the transcription start site (TSS200 or TSS1500), 5’ untranslated regions (UTR) and the first exon; (2) “GeneBody” probes for those located within gene transcribed regions (except the first exon) and 3’UTRs; (3) The remaining probes were defined as “Intergenic” probes.

### RNA extraction and RNA-seq data generation

Total RNA was extracted from keloid tissues or primary fibroblasts using RNeasy Fibrous Tissue Mini Kit (QIAGEN, Cat#74704) and RNeasy Mini Kit (QIAGEN, Cat# 74104) followed by on-column DNase I digestion according to the user manual. Sequencing libraries were prepared using KAPA RNA HyperPrep Kits (Roche, Cat# KK8541) and sequenced on an NovaSeq X or NovaSeq 6000 (Illumina) as 150 bp paired-end (PE150) reads. About 40 million reads were generated for each sample.

### RNA-seq data analysis

The quality control of the RNA-sequencing data was conducted using FastQC. The RNA-sequencing reads were trimmed to remove adapter sequences using Trimmomatic and aligned to the hg38 genome using STAR aligner with proper capture of RNA strand orientation. Then the aligned reads were quantified using htseq-count with GENCODE v37 annotation. Gene counts were then further processed with DESeq2 to identify differentially expressed.

### Cell Culture and Decitabine Treatment

KFs were isolated from freshly excised human keloid tissues as previously established protocols^22,23^. Cells were maintained in Dulbecco’s Modified Eagle Medium (DMEM; Gibco, Cat# 11965092) supplemented with 10% fetal bovine serum (FBS; Gibco, Cat# 10099141C) and 1% penicillin-streptomycin (Gibco, Cat# 15140122) at 37 °C in a humidified atmosphere containing 5% CO₂.

Experiments were conducted using cells between passages 3-7 to ensure optimal viability and consistency. Decitabine (DAC; MedChemExpress, Cat# HY-A0004) was freshly reconstituted in ultrapure water and then sterilized by filtration through a 0.22 μm filter. This reconstituted DAC solution was further diluted in complete medium to prepare final working concentrations of 1, 2, 4, 8, and 16 μM. Given that DAC exerts its mechanism of action during the DNA division phase of cells, the culture medium containing DAC was replenished every 24 hours for the first 48 hours; thereafter, drug treatment was terminated, and cells were cultured without DAC for 5 days before subsequent detection assays were performed. Vehicle control groups received an equal volume of ultrapure water, with the final volume added showing no detectable effect on cell viability or morphology.

### Cell Viability Assay

Cell viability was evaluated using the Cell Counting Kit-8 (CCK-8; APExBIO, Cat# K1018-5) according to the manufacturer’s protocol. Briefly, KFs were seeded in 96-well plates at a density of 3×10^3^ cells/well and treated with indicated concentrations of DAC for 72 h. At each time point, 10 μL of CCK-8 reagent was added to each well, and the plates were incubated at 37 °C for 2 h. Absorbance was measured at 450 nm using a Varioskan LUX microplate reader (Thermo Fisher Scientific). Relative cell viability was calculated as (absorbance of treatment group/absorbance of vehicle control group) × 100%. Each treatment group included six technical replicates, and the experiment was independently repeated three times.

### Cell Proliferation Assay

5-Ethynyl-2’-deoxyuridine (EdU) incorporation assays were performed using the EdU Apollo®567 In Vitro Imaging Kit (RiboBio, Cat# C10310-1) to assess cell proliferation. KFs were seeded in 24-well plates at 2×10^4^ cells/well, adhered overnight, and treated with DAC for 72 h. Cells were then incubated with 50 μM EdU for 6 h, fixed with 4% paraformaldehyde (PFA; Biosharp, Cat# BL539A) for 30 min at room temperature, and permeabilized with 0.5% Triton X-100 in PBS for 15 min. Click reaction was performed using Apollo®567 staining solution for 30 min in the dark, followed by nuclear counterstaining with 4’,6-diamidino-2-phenylindole (DAPI; Beyotime, Cat# P0131) for 10 min. Images were captured by Nikon A1R confocal microscopy, and at least five random fields per well were analyzed. The percentage of EdU-positive cells was calculated as (number of EdU-positive nuclei / total number of DAPI-stained nuclei) ×100% using ImageJ software. Each experiment was performed in triplicate.

### Cell Migration Assays

Following 48h of pretreatment with 2 μM DAC, KFs were trypsinized, resuspended in serum-free DMEM, and seeded into the upper chambers of 24-well Transwell inserts (8 μm pore polycarbonate membrane; Corning, Cat#3422) at 5×10⁴ cells/well. Complete DMEM containing 10% FBS was added to the lower chambers as a chemoattractant. After 24 h of incubation at 37 °C, non-migrated cells on the upper surface of the membrane were removed with a sterile cotton swab. Migrated cells on the lower surface were fixed with 4% PFA for 20 min, stained with 0.1% crystal violet (Solarbio, Cat# C8470) for 30 min, and rinsed with PBS to remove excess stain. Imaged under a light microscope and migrated cells were counted in five random fields per insert using ImageJ. Experiments were independently repeated three times.

### Wound-Healing Assay

KFs were seeded in 6-well plates and grown to 100% confluence. A straight scratch was created in the monolayer using a sterile 200 μL pipette tip, and floating cells were removed by washing twice with PBS.

Cells were then incubated in serum-free DMEM containing 2 μM DAC. Wound closure was monitored at 0 h and 24 h. Wound areas were quantified using ImageJ. The migration rate was calculated as [(initial wound area - remaining wound area) / initial wound area] ×100%. Each treatment group included three biological replicates.

### Western Blotting Analysis

Total cellular proteins were extracted using RIPA lysis buffer (Beyotime, Cat# no. P0013B) supplemented with 1× protease and phosphatase inhibitor cocktail (Beyotime, Cat# P1046) to prevent protein degradation. Protein concentration was determined using the BCA Protein Assay Kit (Beyotime, Cat# P0010) to ensure equal loading. Proteins were separated by 4-20% gradient SDS-PAGE gels (ACE, Cat# ET15420Gel) and transferred onto 0.45 μm PVDF membranes (Millipore, Cat# IPVH00010) at 300 mA for 90 min. Membranes were blocked with 5% non-fat milk in TBST (20 mM Tris-HCl, 150 mM NaCl, 0.1% Tween-20, pH 7.4) for 1 h at room temperature, followed by overnight incubation at 4°C with primary antibodies against COL1A1 (1:2000, Abcam, Cat# AB270993), COL3A1 (1:2000, Abcam, Cat# AB184993), COMP (1:1000, Abcam, Cat# AB74524), POSTN (1:1000, Abcam, Cat# AB14041), BMP2 (1:1000, Abcam, Cat# ab96826), α-SMA (1:1000, Abcam, Cat# AB124964), and β-actin (1:5000, Ray Antibody, Cat# RM2001). After primary antibody incubation, membranes were washed 3 times with TBST (5 minutes each) and then incubated with HRP-conjugated secondary antibodies (1:5000, Ray Antibody, Cat# RM2001; RM3002) for 1 hour at room temperature. Following secondary antibody incubation, membranes were washed again 3 times with TBST (5 minutes each). Protein bands were visualized using Super ECL Detection Reagent (Yeasen, Cat# 36208ES76) and quantified with ImageJ software. Relative expression was normalized to β-actin. All experiments were independently repeated three times.

### Patient-Derived Xenograft (PDX) Model and Treatment

All animal experiments were approved by the Institutional Animal Care and Use Committee of Dermatology Hospital of Southern Medical University and conducted in accordance with the Guide for the Care and Use of Laboratory Animals. Female BALB/c nude mice (7 weeks old, SPF grade) were purchased from Zhuhai BesTest Bio-Tech Co., Ltd. and housed under specific pathogen-free conditions (temperature: 22±2 °C, humidity: 55±5%, 12-hour light/dark cycle) with free access to food and water. Human keloid tissues were obtained from patients who underwent keloid resection at Dermatology Hospital of Southern Medical University, with written informed consent obtained from all participants.

Keloid fragments (5×5×5 mm^3^) were subcutaneously implanted into the dorsal flanks of nude mice (one fragment per mouse). Seven days after implantation, mice were randomly divided into two groups (n=5 per group): DAC treatment group: intra-tumoral injection of 100 μL DAC solution (0.2 mg/mL in 0.9% saline); (2) vehicle control group: intratumoral injection of 100 μL vehicle solution (0.9% saline).

Injections were administered every other day for 14 doses. On day 28, mice were euthanized, and xenografts were excised, weighed, and either preserved in RNAlater™ Stabilization Solution for RNA extraction or fixed in 4% paraformaldehyde for histological analysis.

### Histological and Imaging Analysis

Paraformaldehyde-fixed xenograft tissues were embedded in paraffin, sectioned into 5 μm-thick slices, and mounted on glass slides (CITOTEST, Cat# No. 188105W). Hematoxylin-eosin (H&E) staining was performed to evaluate general tissue morphology. Masson’s trichrome staining was conducted using a Masson Staining Kit (Baso, Cat# No. BA4079A) to assess collagen deposition, and Sirius Red staining was performed using a Sirius Red Staining Kit (Baso, Cat# No. BA4356) to differentiate collagen types. Sirius Red-stained sections were observed under a polarization microscope to analyze collagen birefringence: type I collagen appeared as orange-red fibers, and type III collagen as yellow-green fibers.

Images were captured at 200× magnification. For immunofluorescence staining, deparaffinized and rehydrated sections were subjected to antigen retrieval (pH 9.0, Gene Tech, Cat# No. GT102410), then blocked with 5% normal goat serum (NGS, BOSTER, Cat# No. AB0009) for 1 h at room temperature. Sections were incubated with rabbit anti-POSTN primary antibody (1:500, Abcam, Cat# No. AB215119) overnight at 4 °C, followed by Alexa Fluor® 488-conjugated Goat Anti-Mouse IgG H&L (1:500, Abcam, Cat# No. ab150113) for 1 h in the dark; nuclei were counterstained with DAPI for 10 min. Images were captured at 200× magnification using a Nikon A1R confocal microscope. Five random fields per section were analyzed, with three sections per xenograft.

### RT-qPCR Analysis

After RNA extraction, reverse transcription of 1 μg total RNA was performed using PrimeScript RT reagent Kit (Takara, RR037A) with random hexamers (37°C, 15 min; 85°C, 5 sec).

For qPCR analysis, reactions (10 μL) contained: 5 μL 2× ChamQ Universal SYBR Master Mix (Vazyme, R433-01), 0.4 μL each primer (10 μM) (Table 2), 2 μL cDNA, and 2.2 μL nuclease-free water.

**Table 2.**
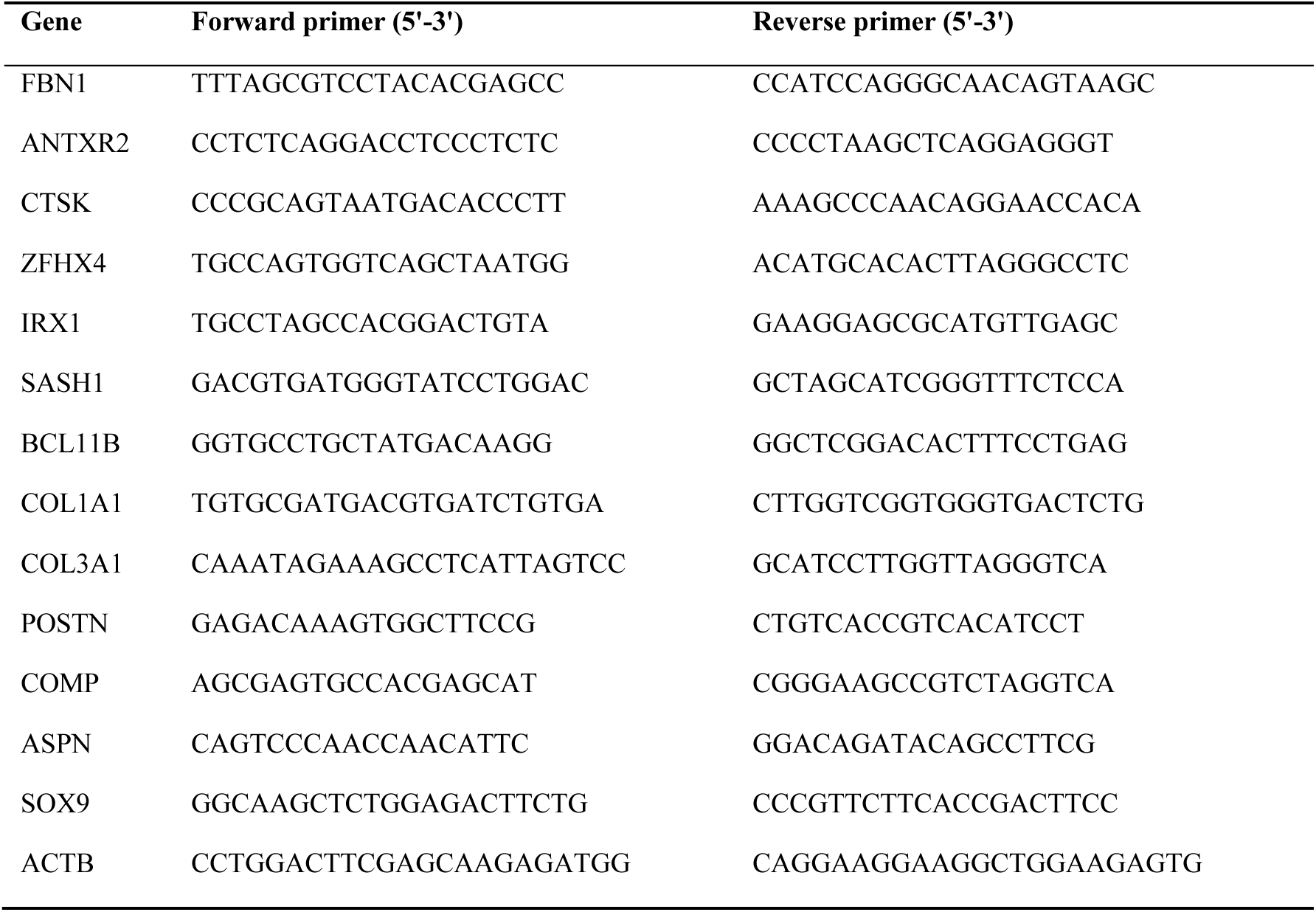
qPCR primer sequences.

Amplification on a CFX Maestro Real-Time PCR System (Bio-Rad) included: 95°C for 30 sec; 40 cycles of 95°C for 10 sec and 60°C for 30 sec; followed by melt curve analysis (60-95°C). All reactions were run in triplicates with appropriate controls.

### Data availability

The previously published DNA methylation data of keloid and normal skin used in this study are available at the GEO database under accession of GSE137134 and GSE56420. Another set of DNA methylation data of keloid are generously shared by Dr. Andrew Stevenson from the University of Western Australia. The data generated in this study are available from the corresponding author upon reasonable request.

### Statistical analysis

Data from cell-based assays and PDX experiments are presented as mean ± standard deviation (SD), with n denoting independent biological replicates. Statistical comparisons were performed using two-sided Student’s t-test for two-group analyses or one-way analysis of variance (ANOVA) followed by Tukey’s post hoc test for multiple comparisons.

For genome-wide analyses, differential expression and DNA methylation were assessed using two-sided Welch’s t-test (matrixTests package). RNA-seq data were standardized using Z-score transformation.

Protein–protein interaction networks were constructed using the STRING database (confidence score ≥ 0.7), and hub genes were identified in Cytoscape using the CytoHubba plugin, ranked by MCC, EPC, Degree, and Closeness algorithms.

All statistical analyses and data visualization were conducted using R (v4.4.1) and GraphPad Prism (v9). A P value < 0.05 was considered statistically significant.

## Results

### Keloid Fibroblasts Retain the Characteristic DNA Hypermethylation Profile of Keloid Tissue

Although primary fibroblasts was a model widely used in keloid mechanism study, it is not clear how this semi-in vivo model can mimic the molecular signatures of keloid tissue. To determine whether primary keloid fibroblasts (KF) retain the DNA methylation features of their tissue of origin, we performed genome-wide methylation profiling on 14 keloid tissues (KT), 4 normal skin tissues (NT), 7 KF, and 3 normal fibroblasts (NF) using Illumina EPIC v2 BeadArray, which was designed to detect methylation level of over 935,000 sites (probes) genome-widely distributed^22^. After filtering the DNA methylation data by removing the data of probes that are (1) linked to known single nucleotide polymorphism, (2) located on the X and Y-chromosomes, and (3) related to age, the data from 785,116 probes were retained for further analysis (Supplementary Figure 1).

First, we characterized the DNA methylation signature of keloid tissues. Compared with NT, 46,643 differentially methylated probes (DMPs) with absolute DNA methylation change (|Δβ|) greater than 0.2 and p < 0.05 were identified in KT, predominantly hypermethylated at 34,637 probes (Figure 1A).

**Figure 1.**
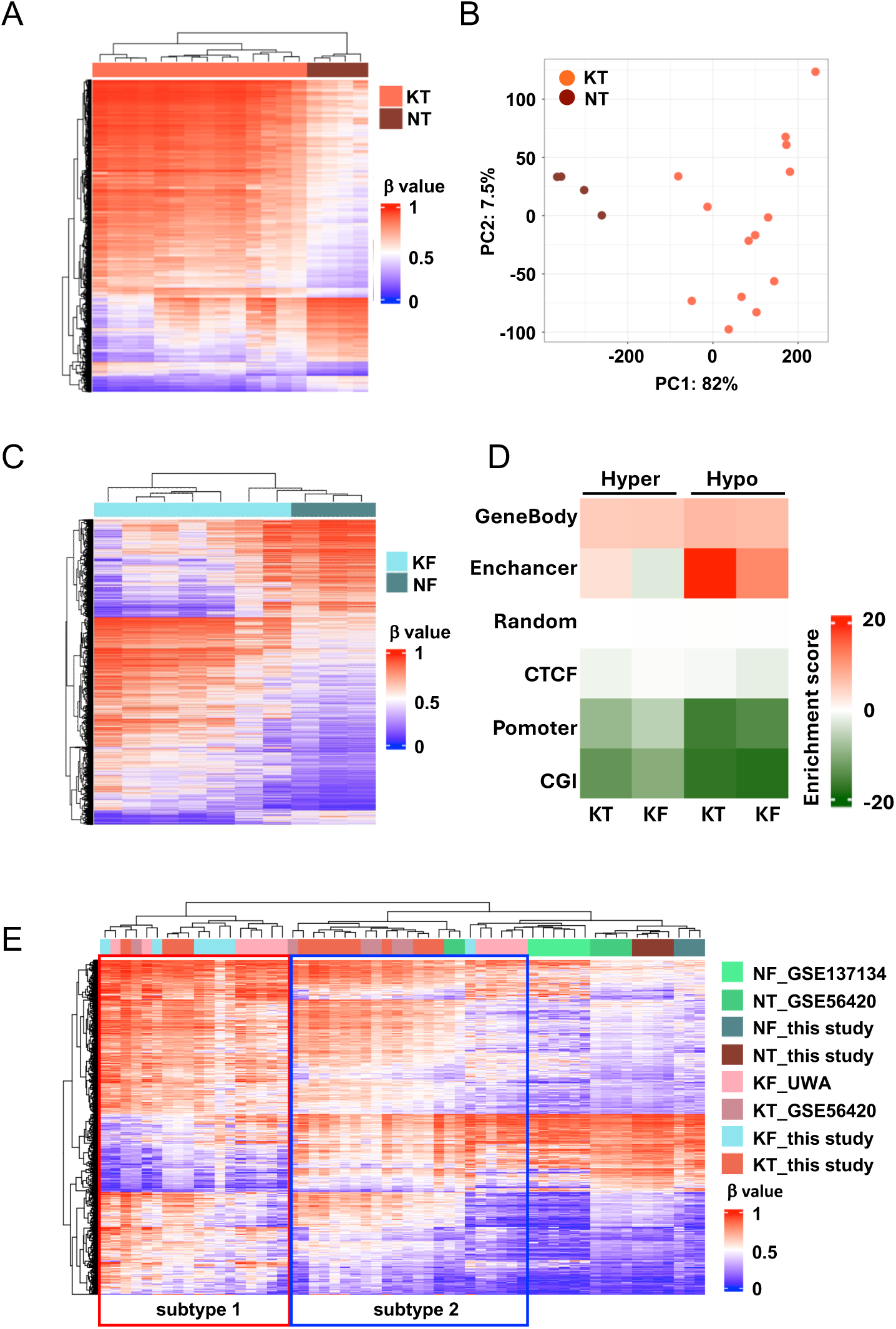
Keloid fibroblasts retain the characteristic DNA hypermethylation landscape of keloid tissue. **(A)** Unsupervised hierarchical clustering heatmap of genome-wide DNA methylation (β value) in keloid tissue (KT; n=14) versus normal tissue (NT; n=4). β value ranges from 0 (unmethylated, blue) to 1 (fully methylated, red). **(B)** Principal component analysis (PCA) of methylation profiles showing KTs cluster separately with NTs. **(C)** Unsupervised hierarchical clustering heatmap of DNA methylation β-values in keloid-derived fibroblasts (KF; n=8) versus normal fibroblasts (NF; n=3). **(D)** Heatmap indicating the enrichment of the DMPs in keloid tissues and fibroblasts at different genomic elements. GeneBody, gene transcribed region (except 1^st^ exon) and 3’ UTR region. Promoter, transcription start site (TSS), 5’ untranslated regions (UTR) and the first exon. CTCF, CTCF binding sites. Random, randomly distributed probes. CGI, CpG island. **(E)** Unsupervised hierarchical clustering showing the cross-cohort validation with three independent public datasets (GSE56420 KT/NT, GSE137134 NF, UWA KF) using 674 shared probes. All keloid samples segregate into two epigenetically distinct subtypes (subtype 1 and subtype 2) that are reproducible across cohorts and platforms.

Principal component analysis (PCA) using these DMPs cleanly separated KT from NT and highlighted marked inter-lesional heterogeneity within the keloid group (Figure 1B). In parallel, KF displayed 12,147 DMPs compared to NF with majority of hypermethylation at 7,697 probes, which is consistent with the finding in keloid tissues (Figure 1C; Supplementary Figure 2A). Cross-comparison identified 3,477 shared DMPs (Supplementary Figure 2A, 1902 hyper- and 1575 hypo-methylated probes) between KT and KF, confirming partial preservation of tissue-level methylation signatures in cultured fibroblasts (Supplementary Figure 2B). We further investigated the genomic regions where the DMPs are located by calculating the enrichment of the DMPs in certain designated genomic elements against their overall distribution in the EPICv2 array. Although the numbers of the DMPs are quite different between KT and KF samples, their distribution patterns are highly consistent, with the enrichment at enhancers and gene body (transcribed region) in KT and KF samples and less enriched at CpG island and promoter regions (Figure 1D).

Furthermore, we merged the keloid-specific DMPs with three independent public cohorts of GSE56420 (6 KTs vs 6 NTs), GSE137134 (6 NFs), and the dataset from University of Western Australia (UWA, including 12 KFs), which were generated using Infinium Human Methylation 450K BeadChip, for validation. Unsupervised clustering based on the β values of the 674 shared DMPs demonstrated a clear separation of keloid samples with normal control samples regardless of the origins (Figure 1E). Notably, all keloid samples, regardless of sample type or cohort, segregated into two methylation-defined subtypes, indicating a reproducible epigenetic heterogeneity that may reflect divergent pathogenic mechanisms or disease severity (Figure 1E). Together, these data validate primary keloid fibroblasts as a biologically relevant in vitro model for studying epigenetically driven keloid pathogenesis.

### Keloid Tissue and Fibroblasts Converge on an Aberrant Osteochondrogenic Transcriptional Reprogramming

To define the transcriptional landscape of keloid pathogenesis, we performed total RNA sequencing on keloid and normal tissues. Compared to normal skin, 3,809 genes were upregulated and 2,030 were downregulated in keloid tissues (Figure 2A; Supplementary Figure 3A). Among the most significantly upregulated genes were established osteochondrogenic markers, including *COL1A1*, *COL3A1*, *FBN1*, *SFRP2*, *COMP*, and *POSTN* (Figure 2A). Gene Ontology (GO) analysis of upregulated genes revealed significant enrichment of osteochondrogenic pathways, including cartilage development (GO:0051216), bone development (GO:0060348), and ossification (GO:0001503) pathways (Figure 2B), consistent with prior reports of osteochondrogenic dysregulation in keloids^7,23^. Multiple collagen related pathways were also identified with the upregulated gene set (Supplementary Figure 3B). The upregulation of osteochondrogenic genes, including *CTSK*, *FBN1*, *ANTXR2*, and *ZFHX4*, was validated by RT-qPCR in an independent cohort (Figure 2C). Downregulated genes were primarily enriched in epidermal development terms, suggesting simultaneous suppression of normal epithelial differentiation (Supplementary Figure 3C).

**Figure 2.**
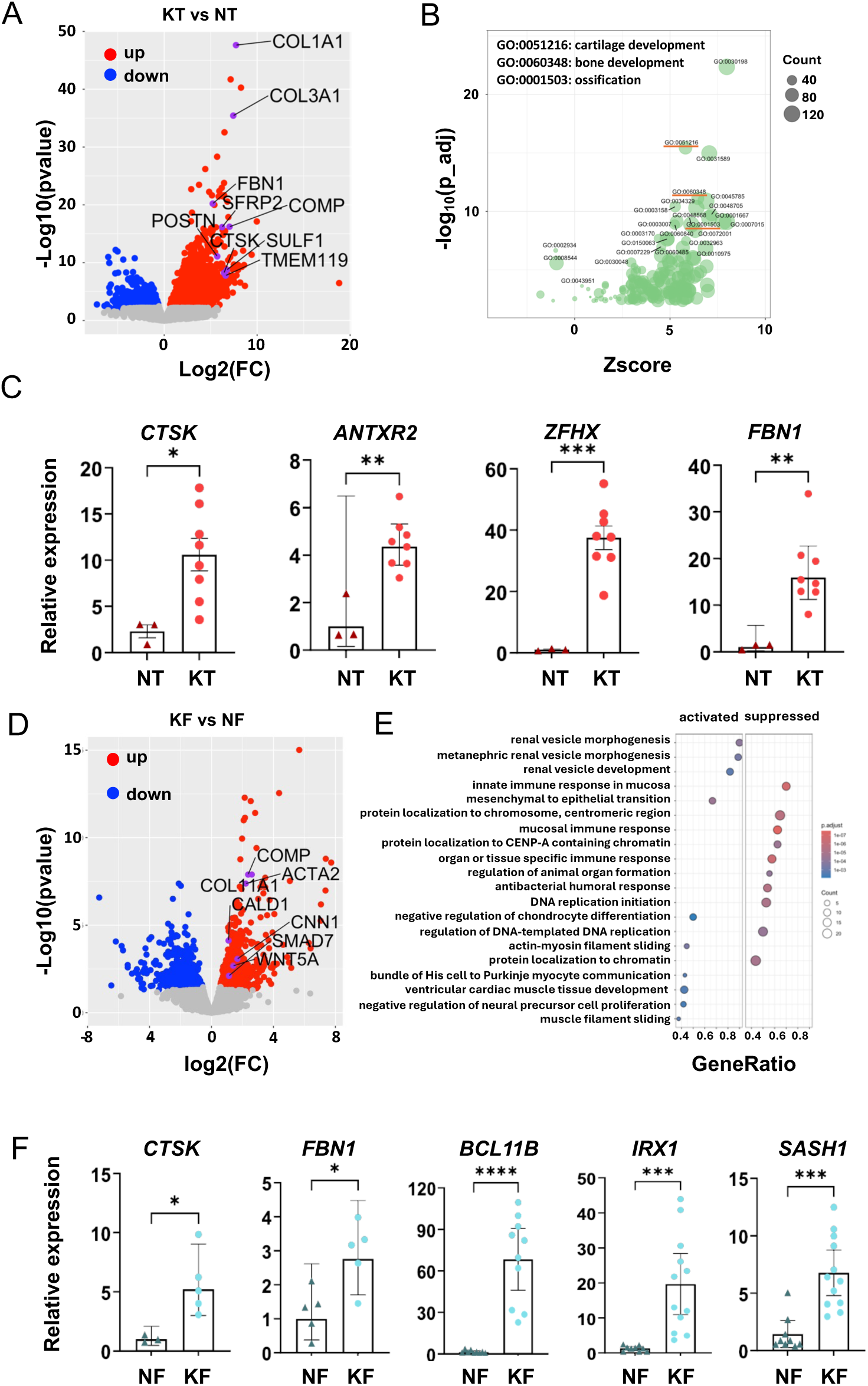
Keloid tissue and fibroblasts converge on an aberrant osteochondrogenic transcriptional reprogramming. (A) Volcano plot of differentially expressed genes in keloid tissue (KT) versus normal tissue (NT); part of osteochondrogenic genes is labelled (*COL1A1*, *COL3A1*, *FBN1*, SFRP2, *COMP*, *POSTN*, *CTSK*, *SULF1*, and *TMEM119*). (B) Gene Ontology analysis for genes upregulated in KT versus NT showing enrichment of osteochondrogenic pathways in biological process term. (C) RT-qPCR validation of the upregulation of osteochondrogenic genes (*CTSK*, *ANTXR2*, *ZFHX4*, and *FBN1*) in independent keloid tissues samples (n=8) versus normal controls (n=3). Data are mean ± SD; *P < 0.05, **P < 0.01, ***P < 0.001, ****P < 0.0001. (D) Volcano plot of the RNA-seq results from keloid-derived fibroblasts (KF) versus normal fibroblasts (NF). Selected osteochondrogenic genes are labelled (*COMP*, *ACTA2*, *COL11A1*, *CALD1*, *CNN1*, *SMAD7*, and *WNT5A*). (E) Gene Ontology of the RNA-seq results of keloid fibroblast. (F) RT-qPCR analysis of the expression of selected osteochondrogenic genes (*CTSK, FBN1*, *BCL11B*, *IRX1*, and *SASH1*) in an independent cohort of fibroblasts from keloid samples (n=8) vs control (n=3). Data are mean ± SD; *P < 0.05, **P < 0.01, ***P < 0.001, ****P < 0.0001.

We next characterized the transcriptome of primary keloid fibroblasts. Compared to normal fibroblasts, 686 genes were upregulated and 616 downregulated in KF, with osteochondrogenic genes, including *ACTA2*, *COMP*, and *COL11A1*, were consistently upregulated (Figure 2D; Supplementary Figure 4A). Downregulated genes in KF were enriched in multiple chromatin and DNA replication associated pathways (Figure 2E), suggesting coordinated epigenetic reprogramming at the transcriptional level. RT-qPCR was performed to confirm the upregulation of *CTSK*, *FBN1*, *BCL11B*, *IRX1*, and *SASH1* in KF versus normal fibroblasts (Figure 2F).

The comparison of tissue and fibroblast transcriptomes revealed 235 shared upregulated genes and 46 shared downregulated genes, confirming partial concordance at the transcriptional level (Supplementary Figure 4B). These findings establish that keloid tissue and keloid fibroblasts converge on a shared osteochondrogenic transcriptional programming, with fibroblasts partially recapitulating the tissue-level phenotype in vitro.

### Methylation Mechanistically Drives the Osteochondrogenic Reprogramming in Keloid Tissue and Fibroblasts

Although aberrant DNA methylation is a common feature of many diseases, only a subset of methylation changes exerts functional effects on gene expression^11,24^. Promoter hypermethylation is associated with transcriptional repression, whereas gene-body hypermethylation can activate transcription, a mechanism recently demonstrated in cancer cells^25^. To determine whether DNA methylation alterations in keloids have functional consequences on transcription regulation, we integrated genome-wide methylation and RNA-seq datasets.

Among the 5,839 DEGs in keloid tissue, 1,492 were accompanied by concordant DNA methylation changes with four regulatory patterns: a, promoter hypomethylation-associated upregulation (n=567); b, gene-body hypermethylation-associated upregulation (n=450); c, gene-body hypomethylation-associated downregulation (n=64); and d, promoter hypermethylation-associated downregulation (n=411) (Figure 3A and B). After removing duplicated genes with multiple probe mappings, 1,362 unique DMP-DEGs (915 upregulated, 447 downregulated) were retained for downstream analysis.

**Figure 3.**
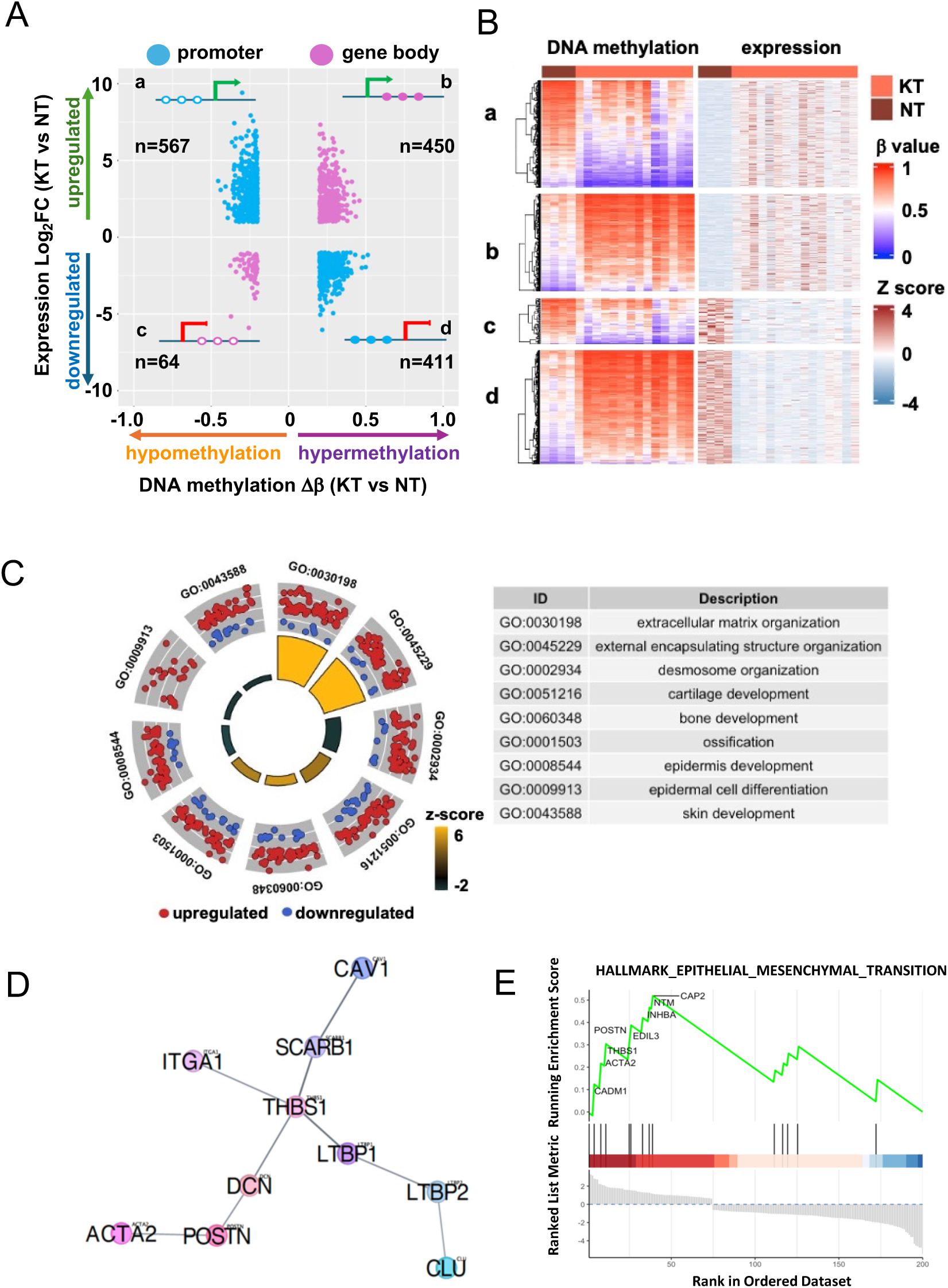
Aberrant DNA methylation mechanistically drives the osteochondrogenic transcriptional reprogramming in keloid. **(A)** Genes regulated by DNA methylation in keloid tissue versus normal skin tissue were plotted with DNA methylation changes at X axis (Δβ, KT vs. NT) and expression change at Y axis (log₂FC, KT vs. NT), Only genes with DMPs located at promoter (blue) and gene body (magenta) were plotted. Groups a–d denote four regulatory classes: a and b, upregulated genes correlated with promoter hypomethylation (a, n=567) and gene-body hypermethylation (b, n=450); c and d, downregulated genes correlated with gene-body hypomethylation (c, n=64) and promoter hypermethylation (d, n=411). **(B)** Unsupervised hierarchical clustering heatmap showing the correlation of gene expression (Z score) and DNA methylation (β value). **(C)** Gene Ontology circle plot showing the enrichment of the pathways, including extracellular matrix organization, cartilage development, bone development, and ossification. **(D)** Protein-protein interaction network of the 201 methylation-associated DEGs in KF, constructed using STRING (confidence ≥ 0.7) and visualized in Cytoscape. Key hub nodes include POSTN, ACTA2, and THBS1, linked to osteochondrogenic differentiation and matrix remodeling. **(F)** Gene Set Enrichment Analysis (GSEA) enrichment plot for the HALLMARK_EPITHELIAL_MESENCHYMAL_TRANSITION gene set in the methylation-linked fibroblast DEG set, driven by POSTN, ACTA2, CAP2, and COMP.

GO analysis of upregulated DMP-DEGs revealed significant enrichment of osteochondrogenic and ossification-related pathways, including cartilage development (GO:0051216), bone development (GO:0060348), and ossification (GO:0001503) (Figure 3C; Supplementary Figure 5A), in keloid tissues. The upregulated cartilage development related genes coupled with both hypomethylation at promoters and hypermethylation at gene body regions (Supplementary Figure 6). Downregulated DMP-DEGs in tissues were primarily enriched in epidermal and skin development pathways (Supplementary Figure 5B), indicating that the aberrant DNA methylation potentially promotes osteochondrogenic reprogramming, as well as suppressing normal epidermal differentiation during keloid formation.

To determine whether this methylation-driven regulatory architecture is recapitulated at the fibroblast level, we performed the same analysis in KF versus NF. By Integrating 12,147 DMPs with 1,284 DEGs, 201 unique DMP-DEGs were identified in keloid fibroblasts (Supplementary Figure 7). Protein-protein interaction analysis of these 201 unique DEGs identified several interconnected networks with key hub nodes including POSTN, ACTA2, and THBS1, all closely linked to osteochondrogenic differentiation and extracellular matrix remodeling (Figure 3D). Consistently, gene set enrichment analysis (GSEA) revealed significant activation of the EPITHELIAL_MESENCHYMAL_TRANSITION hallmark in methylation-linked fibroblast DEGs, driven by osteochondrogenic and mesenchymal genes including *POSTN*, ACTA2, *CAP2*, and *COMP* (Figure 3E). Collectively, these findings demonstrate that DNA methylation orchestrates an osteochondrogenic transcriptional reprogramming in keloids at both the tissue and fibroblast level, providing a mechanistic basis for epigenetically targeted therapeutic strategies.

### Decitabine Suppresses the Osteochondrogenic Phenotype in Keloid Fibroblasts In Vitro

Given the dominant DNA hypermethylation underlies the metabolism reprogramming in keloids, we investigated the therapeutic potential of the DNA demethylating agent (decitabine, DAC) in primary keloid fibroblasts. DAC treatment induced a dose-dependent reduction in cell viability (Figure 4A) and cell proliferation (Figure 4B). At clinically relevant concentrations (1–2 μM), DAC significantly reduced the proportion of EdU-positive cycling cells compared to vehicle control (Figure 4C). We further demonstrated that DAC markedly impaired keloid fibroblast motility by wound-healing assay with DAC treatment for 24 hours (Figure 4D) and transwell migration assay (Figure 4E). Together, these results indicate that DAC suppresses both the proliferative and migratory capacities of keloid fibroblasts.

**Figure 4.**
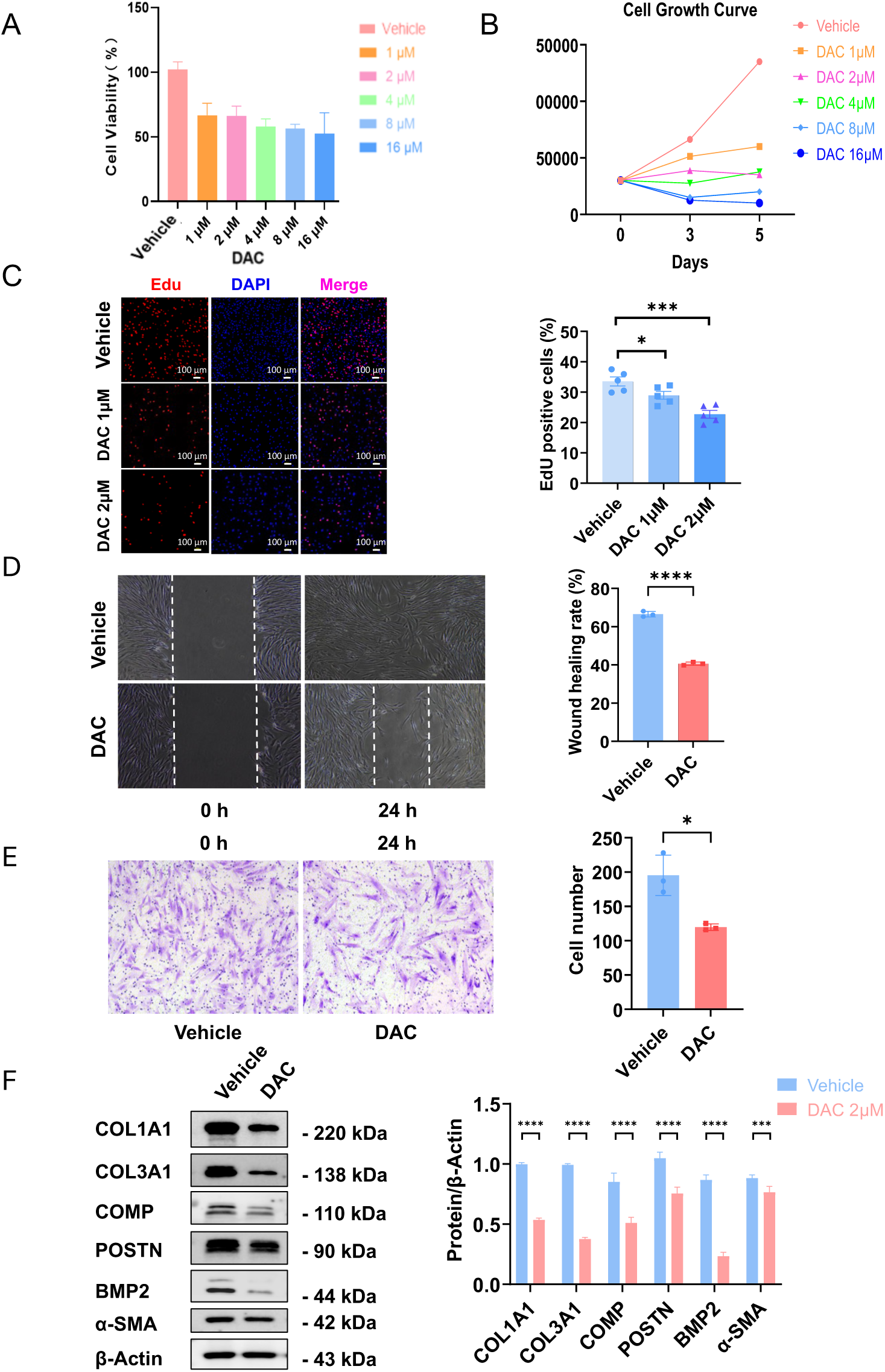
Decitabine (DAC) suppresses proliferation, migration, and the expression of osteochondrogenic markers in keloid fibroblasts in vitro. **(A)** Cell viability of keloid fibroblasts treated with vehicle or DAC (1–16 μM) for 72 hours. Values normalized to vehicle control. **(B)** Cell growth curve for vehicle and DAC-treated fibroblasts in the indicated concentration. **(C)** EdU incorporation assay (red, EdU; blue, DAPI) in vehicle, DAC 1 μM, and DAC 2 μM-treated keloid fibroblasts, with bar graph quantification of EdU-positive fraction. Scale bars: 100 μm. **(D)** Scratch wound-healing assay at 0 h and 24 h in vehicle versus DAC-treated keloid fibroblasts, with quantification of wound healing rate. **(E)** Transwell migration assay at 24 h in vehicle versus DAC-treated keloid fibroblasts, with quantification of migrated cell number. **(F)** Western blot and densitometric quantification of COL1A1, COL3A1, COMP, POSTN, BMP2, and α-SMA, normalized to β-Actin. Data are mean ± SD (n=3–6 independent experiments); ***P < 0.001, ****P < 0.0001.

Western blot analysis revealed the reduced protein level of key osteochondrogenic and fibrotic markers, including COL1A1, COL3A1, COMP, POSTN, and BMP2, as well as the myofibroblast marker α-SMA (Figure 4F), after DAC treatment. These findings indicate that DAC can suppress fibroblasts proliferation by inhibiting osteochondrogenic proteins expression, consistent with the epigenetic mechanism identified in Figure 3.

### Decitabine Suppresses Keloid Growth and Osteochondrogenic Differentiation In Vivo

To evaluate the in vivo therapeutic efficacy of DAC, we employed a keloid patient-derived xenograft (PDX) model. Fresh keloid tissue from consenting patients was implanted subcutaneously in BALB/c nude mice. Animals received intratumoral DAC or vehicle injection every other day for 14 days, and keloid tissues were harvested at Day 28 (Figure 5A). DAC treatment resulted in a significant reduction in tumor weight compared to vehicle-treated controls (Figure 5B), confirming effective suppression of keloid growth in vivo. We observed significant histological changes induced by DAC in the keloid tissues from PDX model. H&E staining demonstrated reduced cellular density and collagen organization in DAC-treated keloids (Figure 5C). Masson’s trichrome staining and quantification of collagen volume fraction (CVF) showed a significant reduction in collagen deposition following DAC treatment (Figures 5D). Picrosirius red staining under polarized light confirmed thinner and less densely packed collagen fibers in DAC-treated tissues compared to vehicle-treated keloids (Figure 5E).

**Figure 5.**
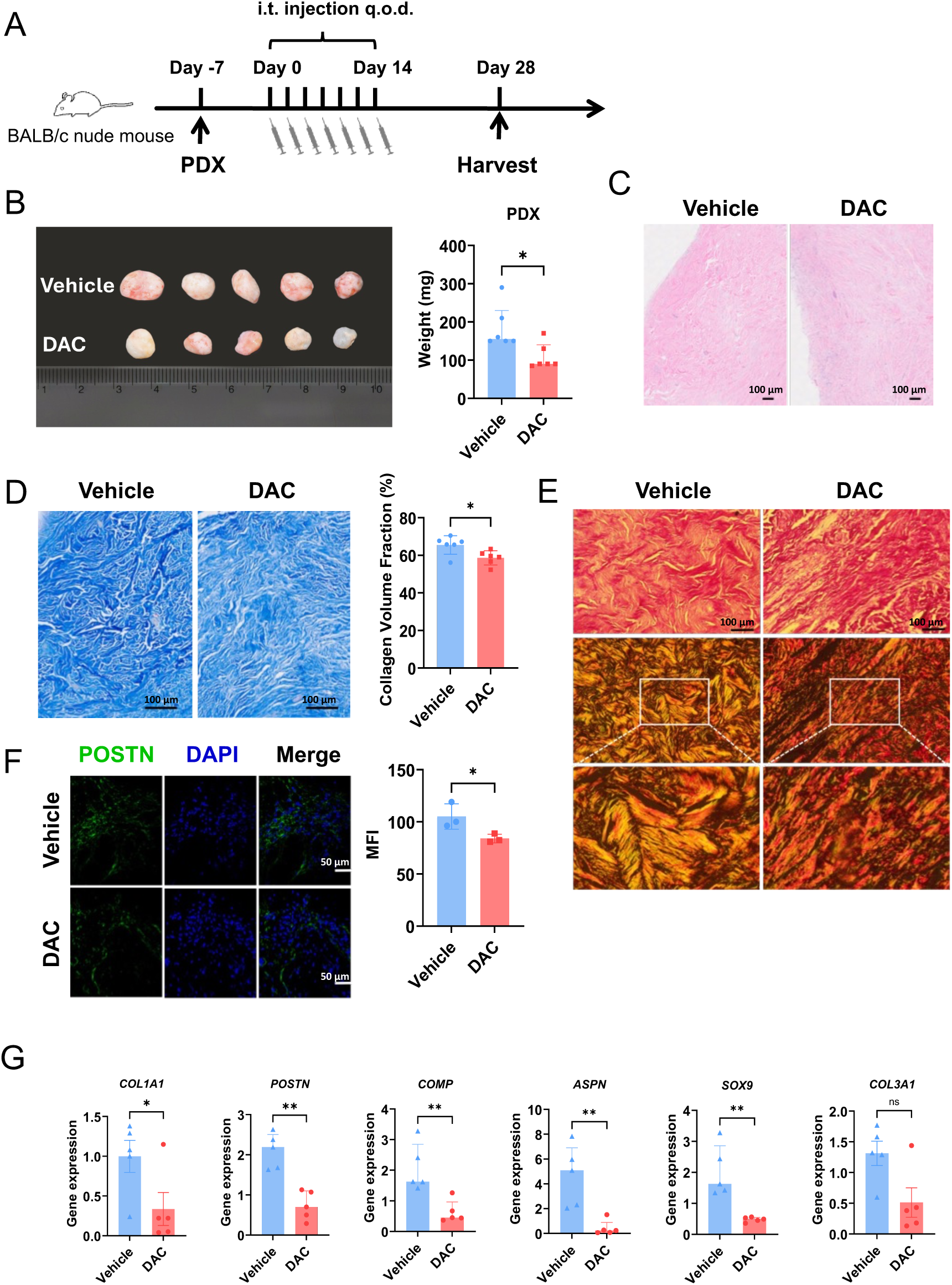
Decitabine suppresses keloid growth and reverses osteochondrogenic differentiation in a patient-derived xenograft (PDX) model in vivo. **(A)** Experimental schematic: keloid tissue was implanted in BALB/c nude mice (Day −7); intratumoral DAC or vehicle injection was administered every other day (Day 0–14); tumors were harvested at Day 28. **(B)** Representative photographs of excised PDX keloids with quantification of tumor weight. n=5 mice/group; *P < 0.05. **(C)** Haematoxylin and eosin (H&E) staining of DAC-treated PDX tissue sections versus vehicle control. Scale bars: 100 μm. **(D)** Masson’s trichrome staining showing collagen distribution in DAC-treated PDX tissue sections versus vehicle control with quantification of collagen volume fraction. n=5 mice/group; *P < 0.05. Scale bars: 100 μm. **(E)** Picrosirius red staining under brightfield and polarized light. Top: low magnification; Middle: polarized light overview; Bottom: polarized light high magnification. Scale bars: 100 μm. **(F)** Immunofluorescence staining for POSTN (green) and DAPI (blue) with quantification showing the decreased fluorescence intensity (MFI) of POSTN immunofluorescence. n=5mice/group; *P < 0.05. Scale bars: 50 μm. **(G)** RT-qPCR showing the upregulation of osteochondrogenic genes (*COL1A1*, *POSTN*, *COMP*, *ASPN,* and *SOX9*) except *COL3A1* in PDX keloid tissue. Data are mean ± SD; *P < 0.05, **P < 0.01.

Next, we detected several known markers by immunofluorescence staining and RT-qPCR. We observed reduced POSTN protein in DAC-treated keloid tissue (Figure 5F) and decreased expression of fibrotic genes (*COL1A1*) and osteochondrogenic differentiation markers (*POSTN*, *COMP*, *ASPN*, and *SOX9*) compared to vehicle, while *COL3A1* showed a non-significant trend towards reduction (Figure 5G). The persistence of *COL3A1* as non-significant is consistent with DAC exerting primary effects on the osteochondrogenic reprograming rather than pan-collagen suppression and warrants further investigation at longer time points.

Collectively, these results provide direct in vivo evidence that DAC suppresses keloid progression by inhibiting the aberrant osteochondrogenic differentiation and extracellular matrix accumulation.

## Discussion

This study establishes DNA hypermethylation as a major, mechanistically coherent driver of aberrant osteochondrogenic reprogramming in keloid fibrosis^7,12,16,23^, and demonstrates that this pathological epigenetic reprograming is reproducible across two independent biological compartments (tissue and fibroblasts), validated in three independent cohorts, and pharmacologically reversible by decitabine both in vitro and in a patient-derived xenograft model. Together, these findings move the field from cataloguing DNA methylation alterations in keloids towards establishing a specific, actionable regulatory mechanism.

Our central mechanistic finding is that gene-body hypermethylation, not only the promoter hypermethylation, play critical roles in driving transcriptional upregulation of osteochondrogenic regulators. This is conceptually consistent with the gene-body methylation-transcription activation relationship demonstrated in cancer by Yang et al^25^, and represents, to our knowledge, the first evidence that this mechanism operates in a fibroproliferative disease context. The key osteochondrogenic transcription factors identified within this aberrant DNA methylation-driven upregulated set, *RUNX2*, *BMP6*, *HOXD3*, *CHST11*, *GHR*, and *TGFBI*, are established regulators of skeletal lineage commitment and cartilage/bone development^26–28^, and their coordinated upregulation via a shared epigenetic mechanism provides a parsimonious explanation for the ectopic osteochondrogenic phenotype that characterizes aggressive keloids.

Although primary fibroblast cultures are widely used in keloid research, their ability to preserve tissue-of-origin epigenetic features has remained uncertain due to epigenetic drift during in vitro expansion^29–31^. We confirmed that KF retain a substantial proportion of the DNA hypermethylation and associated transcriptional signatures observed in KT, and that these disease-specific patterns are reproducible across three independent public cohorts. This multi-cohort validation is a strength of the current study and directly addresses a limitation of prior keloid epigenome studies that relied on single cohorts.

The protein-protein interaction network centered on POSTN, ACTA2, and THBS1, all identified within the methylation-linked fibroblast DEG set, highlights a convergent, matrix-remodeling hub that integrates osteochondrogenic lineage commitment with extracellular matrix assembly. POSTN (periostin) is a secreted matricellular protein strongly associated with fibroblast activation and recurrence in fibrotic diseases^32^; its methylation-dependent upregulation in keloid fibroblasts and its reverse by decitabine in the PDX model make it a candidate pharmacodynamic biomarker for future clinical studies.

The in vivo pharmacologic data from the PDX model are particularly relevant from a translational perspective. Decitabine is already FDA-approved for haematological malignancies; the concentrations effective in our in vitro system (1–2 μM) are within the pharmacokinetic range achievable clinically. The histological reduction in collagen deposition (Masson/Sirius Red) and POSTN immunofluorescence alongside gene expression changes indicates that DAC does not simply suppress fibroblast proliferation but directly reprograms the pathological fibroblast differentiation state. These effects extend beyond anti-proliferative activity and suggest that DAC may prevent recurrence by resetting the epigenetic state driving aberrant osteochondrogenic re-differentiation after excision, a mechanism distinct from, and potentially complementary to, conventional intralesional corticosteroids.

Several limitations should be acknowledged. Cohort sizes, particularly in the methylation analysis, are modest; replication in larger, multi-ethnic cohorts is needed. The PDX model uses intratumoral injection rather than systemic delivery, which may not fully recapitulate clinical pharmacology. Critically, DAC is a global demethylating agent; while our integrative analysis identifies specific methylation-linked gene sets, locus-specific causal validation, for example, CRISPR-based epigenome editing at *RUNX2* or *POSTN*, would strengthen the mechanistic claims and represents an important direction for future work.

Future studies should also incorporate single-cell epigenomic approaches to resolve cell-type-specific contributions to the methylation landscape, and longitudinal clinical sampling to determine whether DAC-induced demethylation is sustained post-treatment.

In summary, we define aberrant DNA methylation as an important epigenetic switch driving aberrant osteochondrogenic reprogramming in keloid fibrosis. We validate primary keloid fibroblasts as a robust model for this epigenetic dysregulation, demonstrate that the regulatory architecture is reproducible across tissue and fibroblast compartments in four independent cohorts, and show that the pathological cell-fate reprogramming is therapeutically reversible by decitabine. By positioning keloids as a paradigm of epigenetically driven fibrotic reprogramming, our findings extend the clinical utility of DNMT inhibitors beyond oncology and provide a mechanistic rationale for prospective evaluation of decitabine-based regimens in keloid management.

## Author Contributions

Bin Yang, Hong-Tao Li, and Chengcheng Deng, and Lian Zhang, Renliang He conceived and designed research. Lian Zhang and Chenmei Liu collected the data; Lian Zhang, Chenmei Liu, Xinyuan Zhou, Hong-Tao Li, Yan Zhang, Ziyan Li, and Shuqing Zhao performed data analysis, prepared figures, and drafted the manuscript. Bin Yang and Chengcheng Deng revised manuscript. All authors have read and approved the manuscript.

## AI Disclosure

During the preparation of this work, the authors used an artificial intelligence tool DeepSeek to assist in the translation and linguistic refinement of the manuscript. The authors reviewed and edited the content as needed and take full responsibility for the content of the publication.

## Acknowledgement

The authors gratefully acknowledge the team of Professor Mark W. Fear and Dr. Andrew Stevenson from the Burn Injury Research Unit at The University of Western Australia for their generous collaboration. We specifically thank them for providing the raw DNA methylation data of keloid fibroblasts. Their willingness to share this invaluable dataset significantly contributed to our research.

